# Integrating computational protein structure predictions and genetic dependencies yields an atlas of human multi-protein complexes (AHMPC)

**DOI:** 10.1101/2025.09.09.675133

**Authors:** Michael Uckelmann, Daniel Alvarez Salmoral, Carmen Maiella, Onno Bleijerveld, Ceri Zwart, Eric Marcus, Joren Brunekreef, Jonas Teuwen, Roderick Beijersbergen, Anastassis Perrakis

## Abstract

Knowledge of which proteins interact to form functional complexes in cells is essential for understanding molecular mechanisms in biology. Structure prediction methods recently allowed to compute the Human Interactome of likely binary protein interactions. We combine computational predictions with orthogonal functional data from the Dependency Map that estimate the correlation between genetic dependencies and vulnerabilities of the corresponding gene pairs. This revealed groups of proteins that likely form larger complexes. Clustering analysis followed by AlphaFold3 multi-protein complex predictions and AlphaBridge analysis provided the basis to construct and atlas of human multi-protein complexes (AHMPC), currently encompassing 354 high-confidence predicted multi-protein complexes. These include well-known assemblies and new ones - such as a complex involving SYS1, JTB, and ARFRP1 that we validate experimentally, suggesting an unexpected role of JTB in Golgi trafficking. To enable the research community to explore the AHMPC and enable further discovery, we cluster all complexes using functional and disease-related embeddings, demonstrate how structured prompts allow validation by large language models (LLMs), and make all structures and analysis available online as an open resource at https://ahmpc.eu/.

## Introduction

Most proteins exert their function through interactions with other proteins and through participation, however transient, in multi-protein complexes. As such, association of a protein with a particular multi-protein complex can provide important information on its physiological function. Dysregulation of protein complexes is oftentimes relevant in the disease context^1^. The structural and functional characterisation of protein-protein interactions is key to understand the function of multi-protein complexes, providing therapeutic opportunities for targeted intervention in the disease context. Various types of protein “glues”^2,3^ and protein binders developed by experimental or computation design^4–10^ are examples for these interventions.

Protein structure prediction has become remarkably reliable with recent computational approaches such as Alphafold3 (AF3)^11^, RosettaFold^12^, D-I-TASSER^13^, CHAI-1^14^, Boltz-1^15^ and others. Increasingly, these models are being applied to predict protein-protein interactions^16,17^. The accuracy of such predictions has now reached levels where ∼18,000 binary protein-protein interactions (PPI) across the human genome have been predicted with reasonable confidence^17^. However, predictions of larger multi-protein complexes remain difficult. The number of possible combinations scales quickly, from ∼200 million for binary interactions among all human proteins, to about a billion for ternary complexes, and trillions for quaternary complexes. This makes it unfeasible to computationally predict all possible multi-protein complexes, and narrowing down the search space becomes necessary.

One way to limit the search space is to augment interaction data with functional data. Individual proteins that together form a multi-protein complex should also show correlated behaviour in functional readouts. The Dependency Map (DepMap) database^18^ provides a rich source of functional information based on gene ablation screens that assess the dependency of hundreds of cell lines on most genes of the human genome for cellular fitness.

Here we combine data from DepMap CRISPR^18,19^ screens with structural predictions of binary protein-protein interactions to focus the search for multi-protein complexes. We identify 354 multi-protein complexes, including well-characterised ones, but also suggesting some that have not been previously characterised. We discuss some of these complexes, and for one we validate interactions, allowing us to propose a role in Golgi trafficking for the jumping translocation breakpoint (JTB) protein, previously shown to play a role in cell division^20^ and mitochondrial function^21^.

To facilitate exploration of the dataset we clustered all complexes according to biological function and association with disease. We also provide structured prompts for large language models (LLMs), describing the supporting data for each complex, enabling further acceleration of the discovery stage. We provide all data as a web-based resource to allow the research community to explore new research directions.

## Results

Μulti-protein complexes can be assumed to represent a coherent structural and functional unit. We would thus expect protein pairs from higher-order complexes to physically interact with each other and contribute to similar phenotypes in cells. As a basis for structural relations we use the recently published predictions of the Human Interactome, an atlas of binary protein-protein interactions across 9,764 unique human proteins^17^ (Interactome dataset in the following). As a proxy for functional similarity we use correlation of gene pairs across the DepMap CRISPR dataset^18^ that measures gene knockout effects on cancer cell survival and proliferation across hundreds of cell lines^19^.

### Creating candidate sets of protein complexes from structural and functional data

We first drew sets of proteins from the interactome dataset^17^ (Fig.1A). For each of the 9,764 unique seed proteins in the interactome set, we identify all direct interactors and all second level interactors of the seed protein, generating candidate sets of potential multi-protein complex members. For each set of candidates, we then construct a matrix and populate it with pairwise correlations across the DepMap CRISPR knockout dataset (Fig.1A,B).

**Figure 1:**
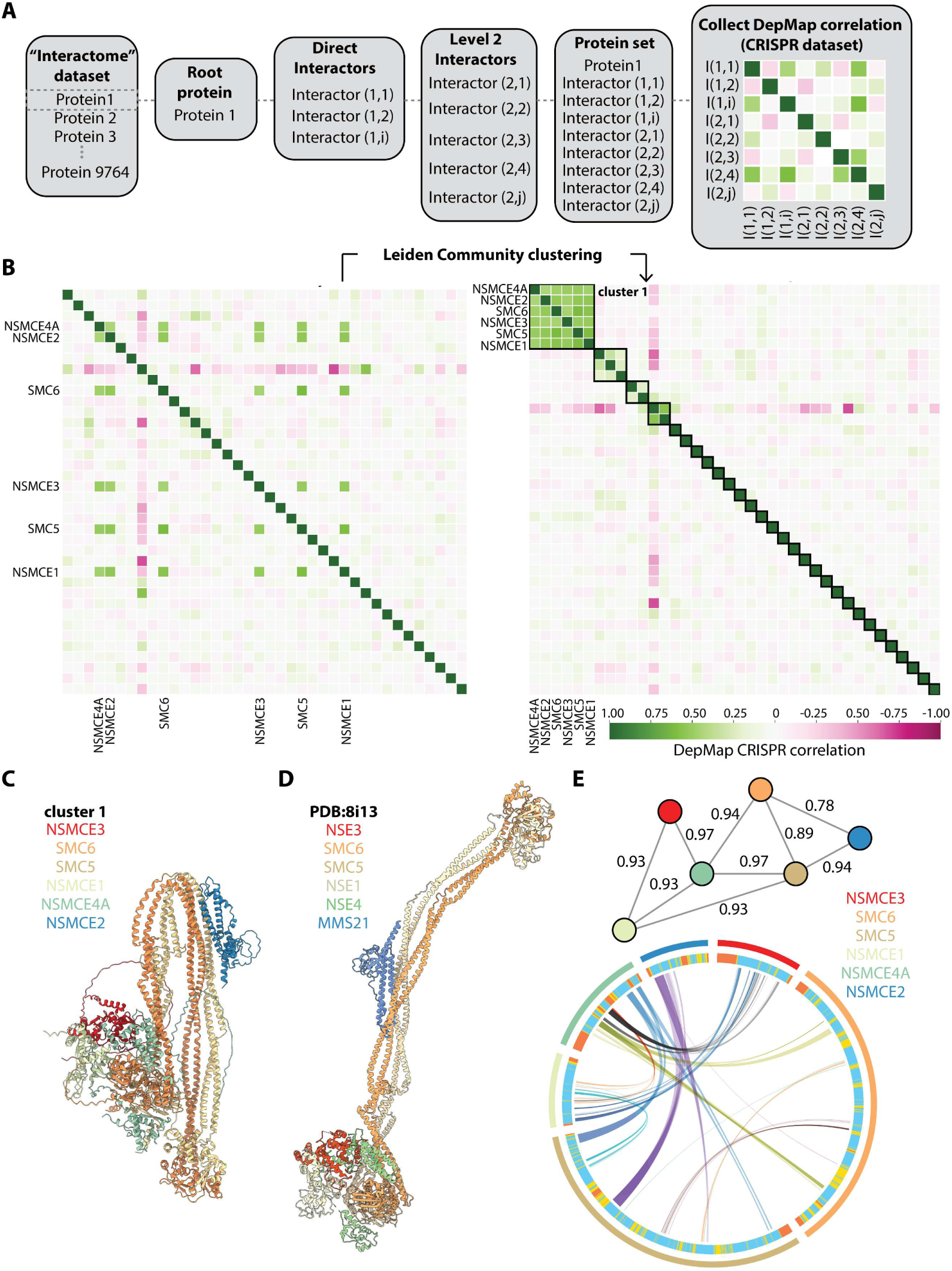
Clustering of orthogonal structural and functional data reveals multi-protein complexes. **A** Flowchart showing how protein sets are drawn and combined in a DepMap correlation matrix. **B** Leiden community clustering of the DepMap correlation matrix generated using NSMCE1 as the root protein. Left: unclustered matrix, right: matrix ordered according to cluster membership. Clusters are indicated by black rectangles. **C** AF3 structure prediction of cluster 1 **D** Experimentally determined structure of yeast SMC5/6 complex (PDB 8i13) **E** AlphaBridge results for cluster 1. In the network representation nodes represent proteins and edges confident interactions. AlphaBridge score is indicated for the edges. In the wheel representation protein chains of are shown as a broken circle with connecting lines indicating confidently predicted interactions as judged by AlphaBridge metrics.

To better illustrate the approach, we show a specific example: combining all direct and second level interactors of seed protein NSMCE1 from the interactome dataset results in a set of 38 unique proteins (Fig.1B). Collecting pairwise DepMap correlations for this set allows to construct the correlation matrix. We then apply the Leiden community detection algorithm^22^, selecting nodes that are more densely connected to each other than to the rest, optimizing for modularity. This reveals several distinct clusters (Fig.1B). “Cluster 1” contains the proteins NSMCE4A, NSMCE2, SMC6, NSMCE3, SMC5 and NSMCE1, which form the well-characterised SMC5/6 complex, involved in DNA repair^23^. The AlphaFold3 prediction for the entire complex shows a conformation of the SMC5/6 complex with the arms folded back, similar to experimentally solved structures of cohesin, condensin and MukBEF^24^ (Fig.1C). Analysis by AlphaBridge^25^ (Fig.1E) highlights many confident interactions across all members of the complex. The experimentally determined structure for yeast SMC5/6 shows an extended conformation with no folding back of the arms (Fig.1D)^26^, suggesting the prediction might show an alternative conformation. The difference might be due to the absence of the Nse2 homologue SLF2 in the structure prediction. Nse2 stabilizes the extended structure of yeast SMC5/6^26^.

This proof-of-concept example shows how leveraging orthogonal structural and functional data can facilitate identification of multi-protein complexes. We applied this pipeline to all 9,764 unique proteins of the interactome dataset. After removing trivial complexes including clusters of histones, keratin clusters, the mito-ribosome, complexes of the electron transport chain and the mediator complex, 712 unique clusters surpassed a mean DepMap correlation cutoff of 0.23.

### Filtering candidate sets by structure prediction and confidence validation

We proceeded to calculate structure predictions for 677 complexes (we removed candidate complexes larger than 5,000 residues due to computational cost) using AlphaFold3, and validated the confidence in the predicted interactions with AlphaBridge^25^. When the AlphaBridge score showed that a member protein of a candidate set was not forming confident interactions with at least one other complex component, that protein was removed. For example, in the cluster containing PEX1, PEX2, PEX6, PEX10, PEX12, PEX26, VPS50 and IFT88, AlphaBridge analysis suggests that VPS50 and IFT88 do not interact with any other member (Supplemental Figure 1A,B). Removing these spurious interactions and re-calculating the AlphaFold3 prediction provides a more confident structure model of the reduced complex (Supplemental Figure 1A-C). Filtering all 677 predictions this way, removed 323 sets entirely (no interactions between any of the proteins). This validation step also reduced 197 sets to less members. We note that some sets include more than one complex: rather than splitting these sets in the separate complexes, we opted to keep them together to show the common origin, and likely related function, of these complexes. As an example, one set includes three different protein complexes (Supplemental Figure 1D, E): the PRC2 core complex (AEBP2, EED, EZH2, SUZ12); a PRC1 RING domain heterodimer (BMI1, RING1); and the MEN1-PSIP1 complex. This is readily apparent from the network representation (Supplemental Figure 1F), where nodes represent proteins and edges confidently predicted interactions. Following the AlphaBridge analysis, we are left with 354 sets (221 with more than two members, and 133 dimers), representing confidently predicted multi-protein complexes, forming the current version of an Atlas of Human Multi-protein Complexes (AHMPC).

### AHMPC and Interactome include some shared but many distinct complexes

The authors of the interactome paper^17^ have suggested their own approach to identify 379 multi-protein complexes from binary predictions, without consideration of functional correlations. They considered clusters of proteins as complexes where each member has at least two interactions with other members. Comparing our suggested complexes (excluding dimers) with this dataset, we find that only two complexes match exactly (Supplemental Figure S2). In 40 cases all the members of an AHMPC complex are contained within a larger Interactome complex. For 18 cases the opposite is the case: a complete Interactome complex is contained in a larger AHMPC complex. A further 120 complexes are partial matches: they have some shared subunits next to distinct additional members. Finally, 41 complexes are completely unique to the AHMPC. Somewhat surprisingly, not all the 133 dimers in the AHMPC are present as confident binary complexes in the Interactome dataset. A total of 55 heterodimers are unique in the AHMPC: FOSB-TGFBR2 or ZSCAN1-ZKSCAN8 are examples of this. There are notable differences between these two resources, despite some overlap. This highlights the added value of combining functional data with the structure predictions.

In summary, the AHMPC provides 354 confidently predicted structures of multi-protein complexes (including dimers). Next to structural information on well-known complexes, the data also suggests the existence of previously unreported complexes, hinting at new biological functions for the proteins involved. Some examples are highlighted in the following.

### Extracting new information from the AHMPC

An important point to consider when interpreting structure predictions for complexes from the AHMPC is that interactions between different subunits are not always predicted with equal confidence. Multiple predictions for the same complex can yield different subunit arrangements in regions where the predicted alignment error is large. As structure predictions need to be interpreted in light of confidence metrics, we use the AlphaBridge scores^25^ to inform users about the confidence in binary interfaces. In light of these considerations, we will present some examples gathered from examining the AHMPC resource.

Complex 22 in the AHMPC, consists of several members from the Peroxin (PEX) family, which is essential for peroxisome biogenesis^27^. PEX3, PEX16 and PEX19 form a well characterised complex, which is important for peroxisome membrane formation and membrane protein import^27^. In addition, the AHMPC suggests that PEX3 can associate with the ACBD5 protein (Fig.2A). The interaction of ACBD5 with PEX3 is the weakest between the complex components (AlphaBridge edge score of 0.77). In fact, multiple predictions of this complex show different configurations but the ACBD5-PEX3 interaction site remains in place (Figure S3). Examining the biology of this complex, we find that ACBD5 is a peroxisomal tail-anchored protein, a class of proteins that insert into the peroxisome membrane via their C-terminal tail^28^. ACBD5 tethers the peroxisome to the ER via interaction with VAPB^29,30^. Proximity ligation results place ACBD5 in the proximity of PEX16 in cells but, to our knowledge, no stable PEX3-PEX16-PEX19-ACBD5 complex has been reported.

**Figure 2:**
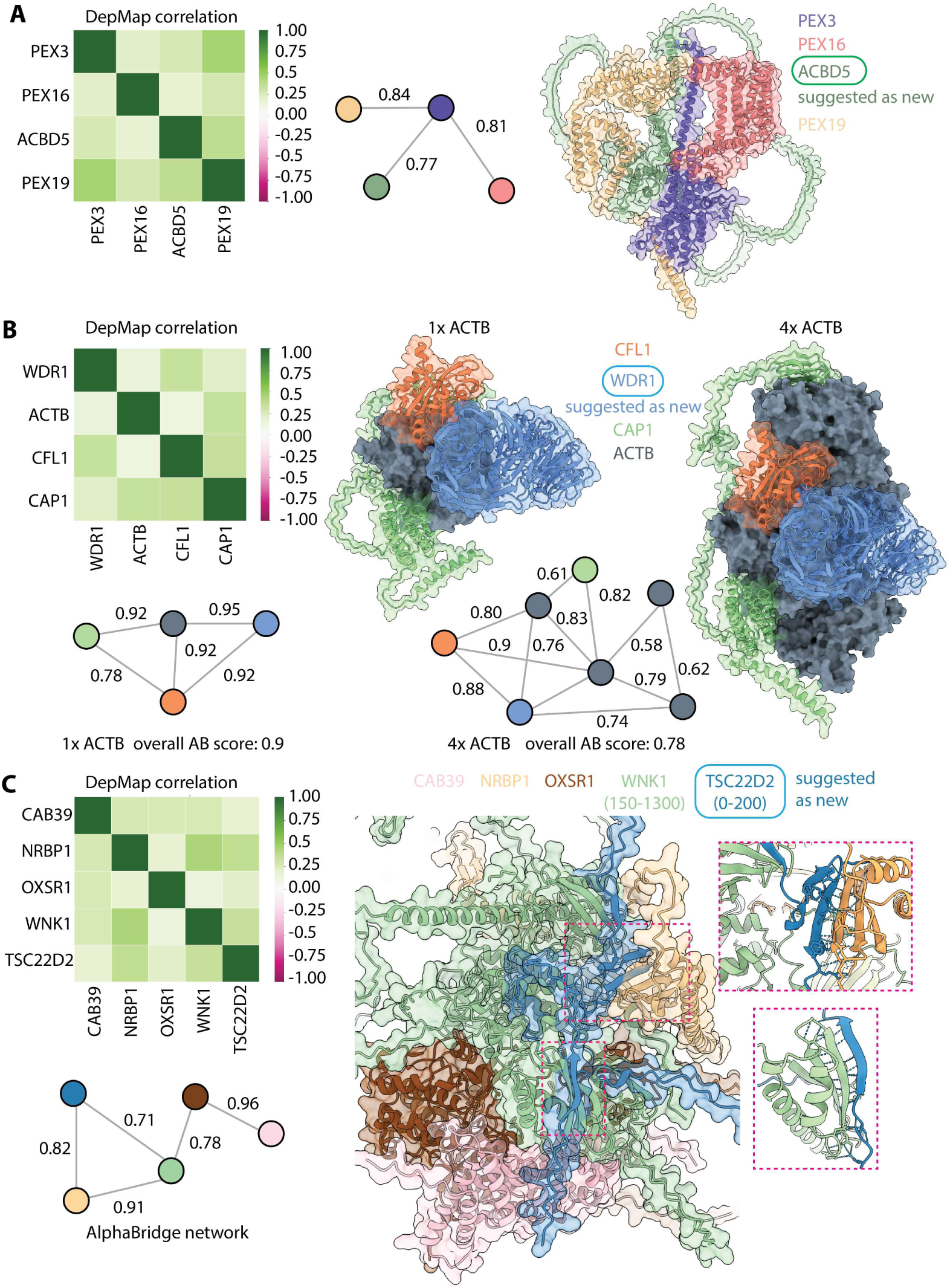
AHMPC suggests new members of multi-protein complexes. **A** DepMap pairwise correlation matrix, AlphaBridge network and AF3 structure prediction for AHMPC complex 22. Gene names of cluster members are indicated. AlphaBridge interface scores are given for each edge. ACBD5 is suggested as a new interactor. **B** DepMap pairwise correlation matrix, AlphaBridge network and AF3 structure prediction for AHMPC complex 294. Gene names of cluster members are indicated. AlphaBridge interface scores are given for each edge. **C** DepMap pairwise correlation matrix, AlphaBridge network and AF3 structure prediction for AHMPC complex 512. Gene names of cluster members are indicated. AlphaBridge interface scores are given for each edge. TSC22D2 is suggested as a new interactor.

Complex 294 includes ACTB (actin), CFL1, WDR1 and CAP1 (Fig.2B). While all these proteins are involved in actin depolymerization^31–33^, a functional complex involving all four was only very recently described (data not published). The predicted structure of the complex bound to monomeric actin shows many confident interactions (Fig.2B). CFL1 directly interacts with both WDR1 and CAP1. CAP1 clamps actin between its N- and C-terminal domains. To see how polymerized actin could affect complex formation, we ran an additional AlphaFold3 prediction including four copies of ACTB (Fig.2B, right panel). While overall the prediction confidence declines for this structure model (AlphaBridge score 0.9 for monomeric ACTB vs 0.78 for four copies), some contacts are still predicted with reasonable confidence. WDR1, binding two laterally associated ACTB molecules, maintains contact with CFL1. The position of CFL1 on the filament reflects an experimentally determined structure^33^. A flexible hinge region in CAP1 allows it to maintain the clamp on the actin filament. We had speculated that the predictions may reflect different states in the actin-severing and recycling cycle^31^, a hypothesis in good agreement with unpublished data from another group.

Complex 512 in the AHMPC includes the proteins CAB39, NRBP1, OXSR1, WNK1 and TSC22D2 (Fig.2C). The relatively low DepMap correlation alone would not necessarily indicate complex formation, but an AlphaBridge score of 0.8 indicates confident interactions are present. This highlights that our pipeline can extract meaningful relations from weak signals. The structure prediction shows that TSC22D2 forms confident interactions with WNK1 and NRBP1 (Fig.2C, disordered regions of WNK1 and NRBP1 have been trimmed for this prediction). TSC22D2 is predicted to interact with both WNK1 and NRBP1 via a β-strand addition (Fig.2C, zoom). AlphaBridge scores of 0.71 and 0.82 on the edges connecting TSC22D2 with WNK1 and NRBP1, respectively, indicate that these interactions are predicted with reasonable confidence. Notably, the regions in TSC22D2 that form the β-strands are predicted to be disordered in the structure prediction of TSC22D2 alone^34,35^, suggesting a disorder to order transition upon complex formation. The inclusion of TSC22D2 in this complex provides a structural rationale for an effect of TSC22D2 on WNK activation, reported in a recent preprint^36^.

### An unexpected role for JTB at the Golgi

Complex 456 in the AHMPC consists of ARFRP1, ARL1 and the single-pass transmembrane proteins SYS and JTB. While the DepMap correlation is moderate, the structure prediction suggests extensive confident interactions among the complex members (Fig.3A,B). ARFRP1, ARL1 and SYS1 are involved in Golgi trafficking^37^. The membrane anchored SYS1 is necessary for ARFRP1 recruitment to the Golgi, where it interacts with ARL1, facilitating downstream signalling^37^ , but mechanistic details are scarce. JTB presents a potential new regulator of GTPase signalling, but has not been previously associated with ARL1 or ARFRP1. In the human interactome^17^, JTB is only recognised as a binary interactor of SYS1, but in AHMPC it also interacts confidently with ARL1 (AlphaBridge score 0.88). JTB’s biological function is poorly characterised, with reports suggesting variable functions in mitosis^20^ or mitochondrial regulation^21^. It is frequently upregulated in cancer^38^ and has most recently been linked to osteosarcoma^39^.

**Figure 3:**
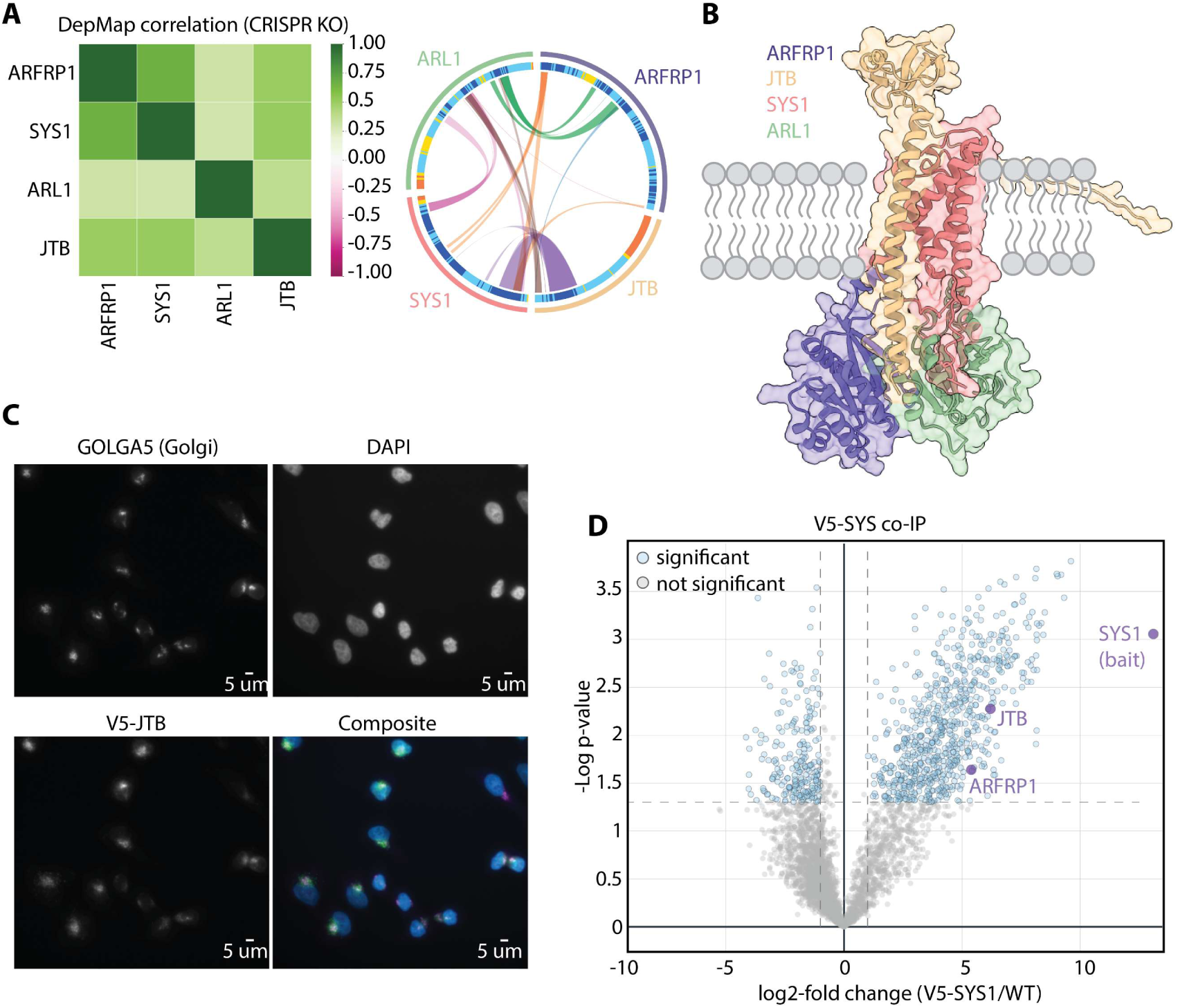
Experimental confirmation of the ARFRP1-SYS-JTB complex. **A** DepMap correlation matrix and AlphaBridge results for the ARFRP1-SYS1-ARL1-JTB complex. Proteins are shown as a broken circle with connecting lines indicating confidently predicted interactions as judged by AlphaBridge metrics **B** AF3 structure prediction of the ARFRP1-SYS1-ARL1-JTB complex coloured by chain. The expected position of the membrane is indicated. **C** Immunofluorescence images of fixed HELA cells expressing V5-JTB. Stained with antibodies against GOLGA5 and the V5 tag. **D** IP-MS using V5-SYS1 expressed in HELA cells as bait. Significance cutoff: Student’s T-test p-value of 0.05 calculated on three replicates

The structure prediction of the complex shows that SYS1 and JTB interact extensively via their transmembrane helices to form a hydrophobic membrane tether (Fig. 3B). The cytoplasmic extensions of SYS1 and JTB form a docking platform for the ARL1-ARFRP1 complex. As this complex was both novel and intriguing, we decided to experimentally validate it in cells.

We introduced V5-tagged constructs of JTB and SYS1 into HELA cells via lentiviral transduction. Interacting proteins were identified using co-immunoprecipitation followed by mass spectrometry (IP-MS) (Fig.3D). Using V5-SYS1 as bait significantly enriched for ARFRP1 and JTB, compared to control cells that have not been transduced (Fig.3D). Using V5-JTB as bait significantly enriched for ARRP1 (Supplemental Fig. S4). Consistently, immunofluorescence images on fixed cells stained with anti-V5 and anti-GOLGA5 (Golgi marker) antibodies showed that JTB colocalizes to the Golgi (Fig.3C). These results, combined with the confident structure predictions strongly suggest the existence of a Golgi-associated complex composed of at least ARFRP1-SYS1-JTB. While ARL1 was not enriched in the MS experiments, it could still transiently interact with the complex in a context-dependent manner, not unexpected for a protein involved in downstream signalling following ARFRP1 interaction^37^. Further analysis of the IP-MS results for V5-JTB shows a statistically significant enrichment of proteins that are annotated as Golgi/ER-associated in Uniprot (Fisher Exact Test on Supplemental Table 1, Supplemental Fig.S4). This further supports the proposed association of JTB with the Golgi. We thus hypothesize an unexpected function of JTB in Golgi trafficking, which warrants further investigation.

### Clustering functional embeddings recaps biological context of complexes

Our initial analysis of the complexes convinced us that the AHMPC is useful for enabling new discoveries by the wider community, and we sought ways to further enable its use. To facilitate exploration of the biological functions we generated embeddings that integrated GO annotations and STRING interaction data^40^ (Supplemental Fig.S5, see methods for details). After dimensionality reduction and computation of cosine similarities, followed by UMAP projection in two dimensions and clustering, distinct functional groupings become apparent (Fig. 4A). This functional profiling revealed that complexes clustered into biologically meaningful groups, reflecting shared molecular roles and pathways. For example, the top GO-terms associated with cluster 1 (Fig.4A and Supplemental Table 2), suggest it includes complexes involved in DNA repair and genome maintenance. Inspecting individual complexes, we indeed find many involved in these processes. Examples include the SMC5-6 complex (Complex 54) involved in genome maintenance^23^ , the MIS12 complex (Complex 430) involved in force-generation at the kinetochore^41^ and the BTR complex (Complex 570), involved in dissolution of Holliday junctions^42^. Cluster 9 includes complexes involved in Golgi biology and vesicle trafficking, including the SYS1-JTB-ARFRP1-ARL1 complex discussed above. Consistently, the complexes discussed previously (Fig.2), fall in clusters reflecting their potential biological function: Complex 22 is a member of cluster 0 (protein stability and degradation; Supplemental Table 2), consistent with known roles for individual members in degradation of peroxisomes^43^; Complex 294 is a member of cluster 8 ( signal transduction and cell migration), consistent with the role of independent complex members in actin depolymerisation; and Complex 512 that includes the kinase WNK1 and associated proteins clusters with proteins involved in protein phosphorylation in cluster 14. Thus, clustering allows to group complexes into common biological pathways, highlighting potential functional interactions between complexes. It can also help to assign function to proteins with unknown biology via their association with specific complexes. We named each cluster by the top GO-term associated with it.

**Figure 4:**
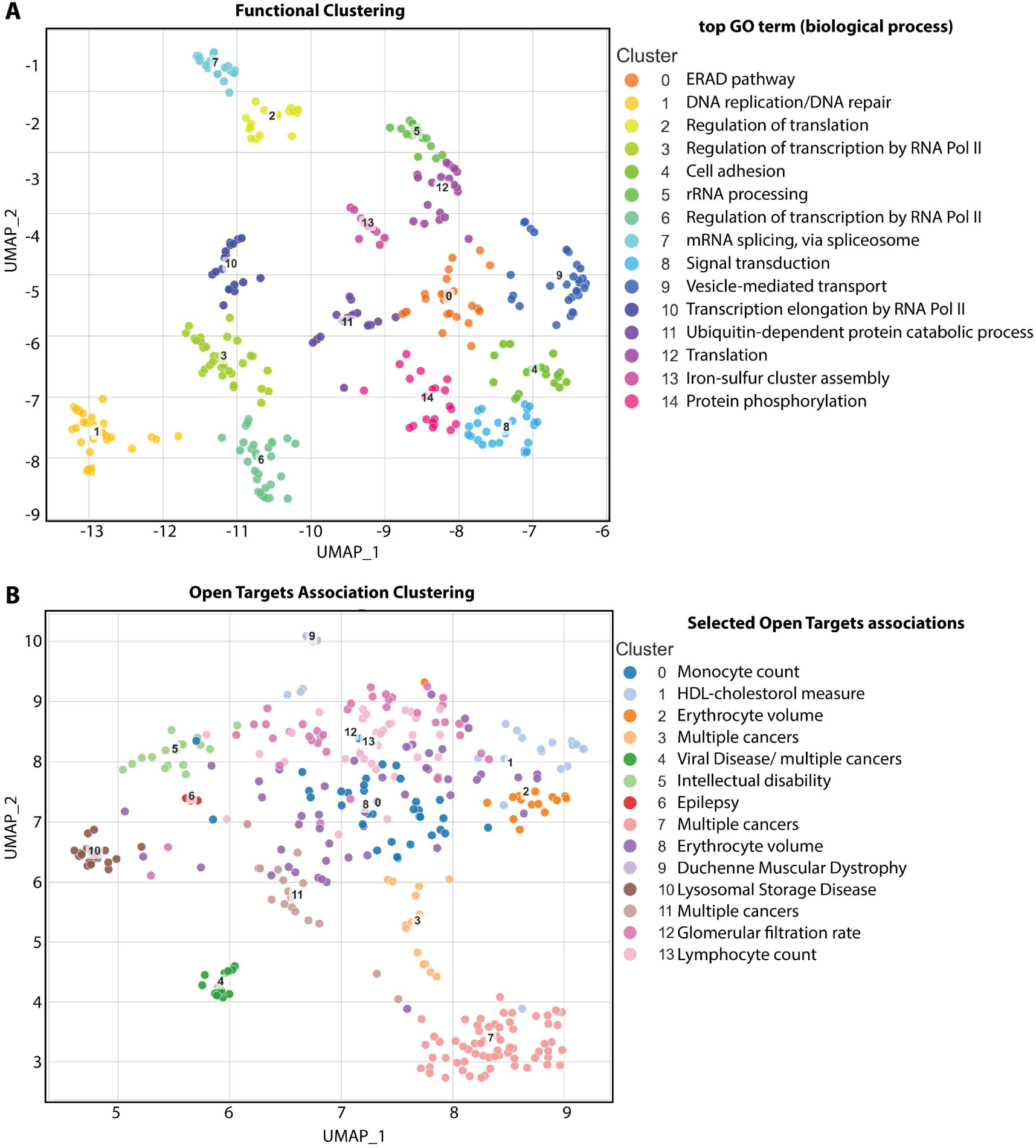
Clustering multi-protein complexes according to biological function and disease association. **A** Clustering based on a functional space defined by GO terms and STRING interactions. Each point represents one complex. The GO-term associated with the most complexes in each cluster is indicated in the legend. See also Supplemental Figure 5 and Supplemental Table 2 for more details. **B** Clustering based on disease association space defined by Open Target scores. Each point represents one complex. One disease association selected from the top 5 associations for each cluster is indicated in the legend. Supplemental Figure 6 and Supplemental Table 3 for details.

To explore the role different complexes may play in a disease context, we followed a similar methodology, making use of data from the Open Targets database^44^. For each complex we collected the Open Targets overall score for all associated diseases, and a feature matrix was constructed (Supplemental Figure 6). Following dimensionality reduction and spectral clustering we obtain meaningful insights into the disease biology of the AHMPC complexes. For example, the top-ranked associations related to cluster 10 suggest that the complexes in this cluster should be involved in neurodegenerative disorders via dysregulation of the endosome-lysosome pathway (Supplemental table 3). This is consistent with the function of individual complexes, many of which are directly or indirectly involved in this pathway: the V1 complex of vacuolar ATPase (Complex 35) involved in acidification of intracellular compartments^45^; the ARP2/3 complex (Complex 12) involved in autophagy and lysosome regeneration^46^; and many spliceosome components that may be indirectly linked. The ARFRP1-SYS1-JTB-ARL1 complex discussed above is also a member of this cluster. It will be interesting to probe the role of JTB and the other members in this disease context. From the previously discussed complexes, Complex 294 and 512 (Fig.2b,c) are members of a large central cluster (cluster 8) that is not very well resolved and associated with haematopoiesis Complex 22 which as we previously discussed is involved in peroxisome biogenesis, a process often dysregulated in neurodevelopmental diseases, is a member of cluster 5 that is associated with intellectual disability. Other examples include complexes involved in viral infections in cluster 4 and complexes involved in haematopoiesis in cluster 18. Clusters 1, 3 and 15 are large clusters that include associations with multiple cancers.

### Creating an opportunity for LLM-accelerated data analysis

Initially, identification of the first complexes that are now in the AHMPC had been followed by extensive, time consuming, manual literature searches. In light of the meaningful functional clustering, we sought new means to accelerate user-oriented analysis of the AHMPC. We opted to explore the use of LLMs to accelerate the initial work involving literature and database searches. To achieve this, we extended our pipeline to output a structured prompt for each complex. This includes the research task (same for all complexes), the top 50 STRING interactions and the DepMap correlation matrix. The prompts for complexes that we had already analysed manually (discussed above), were then submitted to some of the popular current generation (May 2025) LLM platforms.

Based on subjective evaluation of the quality of the outputs, with particular focus on scientific accuracy, we decided to continue with the agentic FutureHouse Falcon model. Falcon takes about ten minutes to generate a detailed report, appropriately referenced, with conclusions similar to the ones we reached after several hours of research for the SYS1-JTB-ARFRP1-ARL1 complex. Falcon places complex 22 in the appropriate context of peroxisome biogenesis and suggests ACBD5 as a novel member. For complex 294, the LLM accurately describes a proposed role in the actin polymerisation cycle. Complex 512 is identified as a WNK-signalling complex, TSC22D2 is suggested as a novel member and a role in ion homeostasis and stress response is hypothesized. In addition to highlighting potentially novel complexes, the AI model also accurately describes the biology of well-characterised multi-protein complexes (e.g. complex 54). Encouraged by our experience, we have constructed structured prompts for all 354 complexes in AHMPC and have executed 76 in Falcon (only selected complexes due to rate limits). We note that the structured prompts can be used with any current or future LLM framework and are available for download from the web resource.

### The AHMPC web resource

The AHMPC is available at https://ahmpc.eu/. All complexes can be accessed from the front page table, which can also be sorted by any of the three quality indicators, and is searchable by any gene or protein identifier (Fig.5A). Clicking a complex, links to an individual summary display page. This shows protein names for each complex member, the DepMap correlation matrix, as well as three representation of the structure of the complex with increasing detail: a network representation including interactions scores for each interaction, the Alpha-Bridge diagram used for validation (Fig.5B), and an interactive 3D presentation of the AF3 structure prediction. More detail on interactions is available by clicking the AlphaBridge link. The structure file for each complex prediction is also available for download. Where available, an AI-generated “Executive Summary” and a collapsible detailed analysis for the complex are populated from the FutureHouse Falcon response to the structured prompt; the structured LLM input prompt is provided for download as a JSON file for every complex. We encourage everyone to use the LLM prompts to test other AI models and work out improvements. We invite the community to scrutinize and use this rich dataset with the hope to stimulate discussion and drive new research directions.

**Figure 5:**
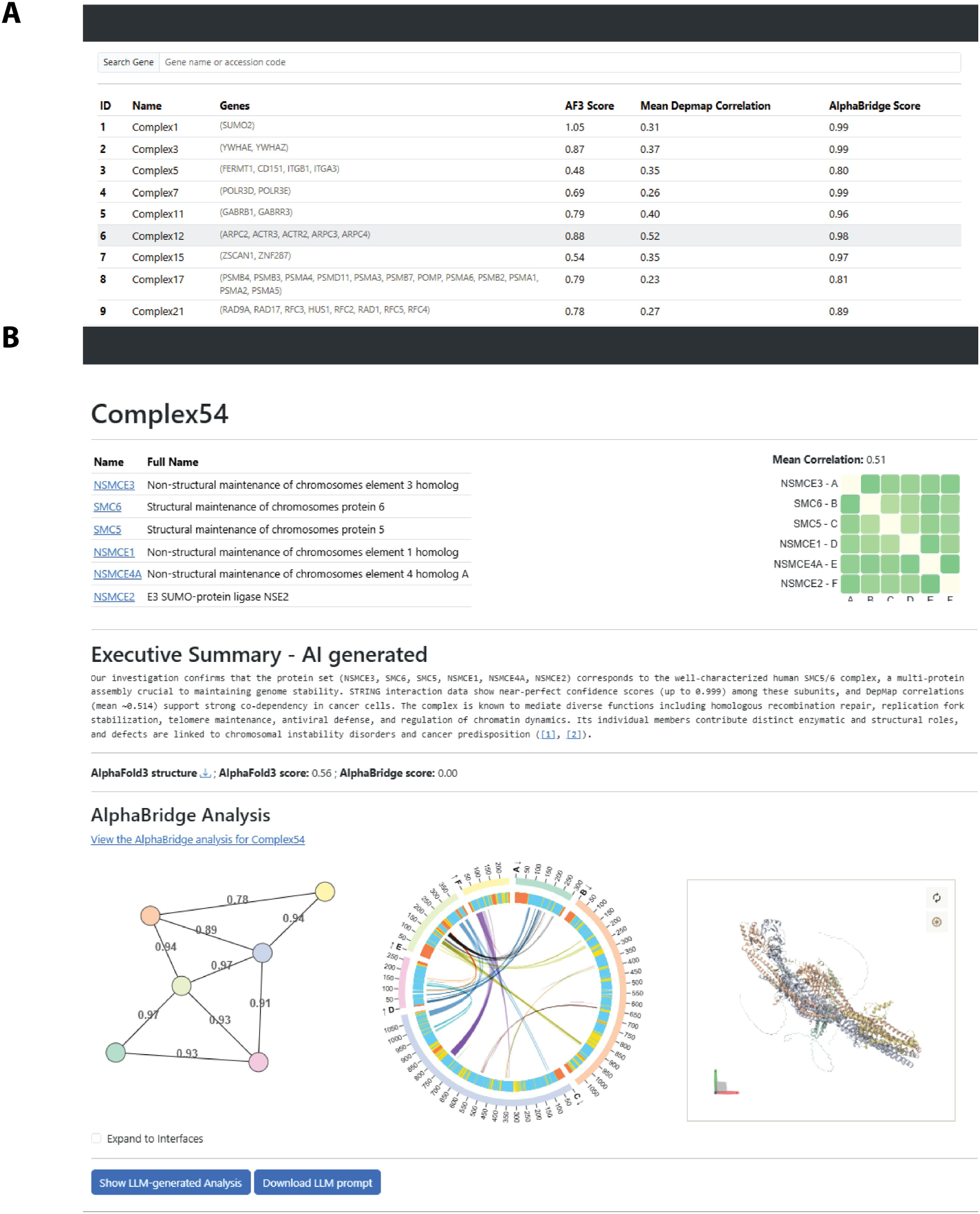
Screenshots from the AHMPC website. **A** The landing page with a table view of the complexes and search function. **B** The display page for complex 54 showing DepMap correlation, Executive Summary, AlphaBridge results as network and detailed depictions and the structure model.

## Discussion

Protein structure prediction methods made incredible advances in the last five years^11–13^ and now provide a remarkable amount of single chain protein structure predictions^35^. Recent efforts have shown the utility of these models for large-scale predictions of binary protein-protein interactions^17^. However, the relevant functional units in which proteins act are often multi-protein complexes. Prediction of these assemblies has remained challenging due to the large number of possible combinations and the scaling of computational time with complex size. Hybrid approaches combining structure predictions with experimental data are now gaining traction as a way tackle this problem^47^. Our study illustrates how combining orthogonal datasets, structural data on binary protein-protein interactions^17^ and DepMap functional screens^18^, can help to identify multi-protein complexes.

The aim of this study is not to provide a complete inventory of human multi-protein complexes. We rather aimed to generate structure-function insight which can lead to biological discoveries. In total, the AHMPC comprises 354 potential protein complexes (including dimers). For each complex we provide the underlying data, as well as validation criteria, interactive inspection tools and, for some cases, LLM-generated annotation. Those data are meant to provide guidance to generate hypotheses that will need to be experimentally verified. They should not be seen as substitutes for thorough experimental characterisation^48^ or literature analysis, but as starting points for accelerated discovery.

An example of such accelerated discovery is the work we present for the role of the JTB protein in Golgi trafficking. Starting from its association with SYS1, ARFRP1 and, potentially, ARL1, we generated experimentally evidence that support the hypothesis that these proteins are associated with each other and the Golgi.

The clustering approaches we presented provides different viewpoints of the AHMPC. Functional annotations of protein structure predictions based on GO-terms and STRING interactions have been implemented for single chain predictions in a hierarchical manner^49^. Here we took a slightly different approach, analysing how multi-protein complexes cluster in a functional space based on weighted GO-term and STRING contribution. Rather than annotating only single proteins, this approach focuses on functional relations between protein complexes. Similarly, GWAS associations have been used to study protein-protein interactions based on network analysis^50^. Here we approach this from a slightly different angle, asking how given multi-protein complexes cluster in a disease space defined by Open Target scores. This may prove useful to identify combinations of multi-protein complexes to target in a specific disease context. Given steadily accumulating data for both structure predictions of protein complexes and functional associations, we expect that similar, perhaps more refined, clustering approaches might help to identify valuable targets for therapeutic interventions.

Finally, our use of a current-generation agentic AI framework, given simple structured prompts, offers a tool that can significantly accelerate exploring the AHMPC for new users. While we caution against over-reliance on AI and want to stress that rigorous scrutiny of the outputs by trained scientist is critical, employing AI frameworks in this way could save a remarkable amount of research hours.

We expect that opening the AHMPC resource to the research community will accelerate new discoveries and might lead to unexpected insight in well-studied individual proteins or protein complexes. Moreover function- or disease-centric browsing of the resource can provide functional associations or lead to the identification of new targets for therapy for specific pathologies.

## Acknowledgements

This work has been supported by an institutional grant of the Dutch Cancer Society and of the Dutch Ministry of Health, Welfare and Sport. A.P. is an Oncode Institute investigator. MU and DAS are (part-time) supported by the Oncode Accelerator project. C.Z. and O.B. are supported by the Dutch NWO X-omics Initiative. We thank the NKI microscopy facility for training and access to light microscopes and the Research High Performance Computing facility of the Netherlands Cancer Institute for providing and maintaining computation resources. We thank Gian-Luca McLelland for providing advice on lentivirus generation and Golgi imaging. Maarten Hekkelman for developing the front end of the AHMPC, and Robbie Joosten for advice on computational matters.

## Methods

### Combining DepMap and Interactome data to find candidate multi-protein complexes

The basis for the DepMap data is the release 24Q2 Post-Chronos CRISPR knockout gene effect estimates (available at https://depmap.org/portal/data_page/?tab=allData “CRISPRGeneEffect.csv”)^18^. Pairwise Pearson correlations were calculated for all combinations across the whole dataset. Sets of proteins that represent candidate multi-protein complexes were drawn from the Interactome dataset^17^. We focussed only on the proteins judged to engage in at least one confident binary interaction with another protein, according to the metrics described in the Interactome manuscript^17^ (set available for download as “GoodPairs.txt” at http://prodata.swmed.edu/humanPPI/bulk_download). We then filtered out those proteins that are not present in the DepMap dataset. For each unique protein in this dataset (9764), we collected all direct interactors and second-level interactors (interactors of interactors), giving us 9764 sets of proteins that may form a multi-protein complex. For each set we constructed a correlation matrix, collecting the pre-calculated pairwise Pearson correlations of the DepMap 24Q2 CRISPR gene effects. We then used the Leiden community detection algorithm^22^ to identify clusters of proteins that show high co-corelation across the DepMap dataset. Graphs were constructed using the igraph library^51^ with proteins as vertices and DepMap correlation as edges. The algorithm was run for 40 different resolution factors in a linear range between 0.01 and 0.2 with the constant Potts model quality function (CPMVertexPartition). For each of the 40 runs, clustering quality was evaluate by calculating the Davies-Bouldin index. The clustering result with the lowest Davies-Bouldin index was used for further analysis. We filtered out clusters that have less than 3 members and those with a mean DepMap correlation of less than 0.23. Further filtering removed clusters of histones and keratin clusters, as well as the mito-ribosome, complexes of the electron transport chain and the mediator complex, because of their large sizes. Redundancy was reduced by merging clusters that are 80% similar (Jaccard similarity) and those where the majority of the set (cutoff 80%) is included in another, larger set. This left a total of 712 candidate complexes. For 677 (excluding complexes larger than 5000 residues) we calculated AlphaFold3^11^ structure predictions using a local AF3 installation.

### AlphaBridge filtering

Each of the structure predictions for the 677 complexes was analysed using AlphaBridge^25^, a tool that determines confidence of interaction sites from AlphaFold3 inputs. We chose an AlphaBridge score cut-of of 0.5 to define confident interactions. Chains from the AF3 prediction that don’t form any confident interactions were removed from the set and a new prediction of the reduced set was calculated. 323 predictions showed no confident contacts between any of the chains and were thus removed entirely, leaving final predictions for 354 complexes.

### Clustering based on GO-terms and STRING interactions

To get a set of GO-terms for each complex, the GO-terms associated with each protein were collected from Uniprot and combined. Each GO-term was only counted once per complex, even if it was associated with multiple individual proteins. Similarly, all STRING interaction partners with the associated combined score were collected for each member of a complex from the STRING interaction database. Where multiple members of a complex interact with the same partner, only the higher score was kept.

A binary GO-term feature vector was generated for each complex where a value of 1 indicates that this GO-term is associated with the complex. The length of the vector is equal to the number of all GO-terms associated with any complex across the whole dataset. A STRING feature vector was generated for each complex in a similar manner. The length of the vector is equal to the number of all STRING interactions across all complexes. For each complex, the STRING interaction score for each interaction associated with this particular complex is included, while the value 0 indicates no interaction with this partner. GO-term feature vectors and STRING feature vectors were combined in two separate matrices of dimensions (# of complexes) x (# of all GO-terms) and (# of complexes) x (# of all STRING interactions), respectively. Both matrices are sparse. Dimensions of both matrices were reduced using truncated singular value decomposition (scikit-learn implementation TruncatedSVD), choosing the dimensions so that >70% of the variance was conserved (65 dimensions for STRING and 100 dimensions for GO matrix). Cosine similarities were then calculated on the lower-dimension matrices. The similarity matrices were then added with a weight of 0.6 applied to the STRING matrix and a weight of 0.4 to the GO-term matrix. Finally, the combined similarity matrix was reduced to two dimensions using UMAP python implementation with the following parameters: umap.UMAP(n_components=final_dim, metric=’precomputed’, random_state=42,init=’random’, n_neighbors=20, min_dist=0.05, n_epochs=10000). Clusters were identified after UMAP using the K-means algorithm with 15 final clusters. 15 clusters were chosen because it maximized the silhouette score. Complex 88 was not included in the clustering (due to late addition).

### Clustering based on Open Targets disease associations

To generate an Open Targets disease association matrix for the complexes, first the protein IDs for each complex were converted to Ensembl IDs. The Open Target disease associations for each complex member were collected using Open Targets API. If multiple proteins of a single complex share a specific disease association we summed the scores.

To reduce noise and exclude features that might not contribute meaningfully to the clustering, we filtered out diseases that had a maximum association score of less than 0.4 across all complexes, a sparsity ratio of >=0.99 and a variance of <=0.01. We also removed features that have particularly high association scores for most of the complexes, because clustering may be dominated by these terms and differences might be masked. The specific associations we removed were: ’neurodegenerative disease’, ’body height’, ’body mass index’, ’body weight’, ’systolic blood pressure’, ’appendicular lean mass’, ’neuroimaging measurement’. The final number of features was 1141.

We then reduced the dimensionality to 65 dimensions using truncated SVD (scikit learn implementation), which maintained >85 % of the variance (Supplemental Figure 4B). After calculating cosine similarities we used spectral clustering (n_clusters =14), with the cosine similarity matrix as input, (scikit learn implementation) to group the complexes. For visualisation, dimensions were reduced to two using umap. Complex88 was not included in the clustering (due to late addition)

### Comparison of Interactome and AHMPC

The Interactome resource^17^ proposes 379 multi-protein complexes. For some of them, protein members can have multiple homologues, of which the authors of the Interactome choose one. To align our choice of homologues with the DepMap data we used the homologues that gave the highest mean DepMap correlation. For most of the 379 complexes we generated matrices of all possible homologue permutations and chose the combination that had the highest mean DepMap correlation. For few large complexes the number of permutations was exceptionally large, for those we randomly chose possible combinations until the total number of matrices reached 10671.

### Lentivirus generation

3.5 x 10^6^ HEK293t cells were seeded into a 10cm culture dish. The following morning, 1.3 pmol of packaging plasmid psPAX2, 0.7 pmol of envelope plasmid pMD2.G and 0.8 pmol of the respective transfer plasmid encoding SYS1 or JTB were combined in opti-MEM in a total volume of 500 ul. 60 ug of PEI were mixed with 500 ul of opti-MEM in a separate tube. The diluted PEI mixture was mixed with the diluted DNA mixture and incubated for 15 minutes at room temperature. After incubation, 10 ml DMEM supplemented with 10% FBS, PenStrep and Glutamine was added, the medium from the 10 cm plates was aspirated and the transfection mix was added. The following morning, the transfection media was aspirated and replace with DMEM supplemented with 10% FBS, PenStrep and Glutamine. Virus was harvested 48 hours later and used for infection of HELA cells directly.

### Transduction of HELA cells

HELA cells were maintained in DMEM supplemented with 10% FBS, PenStrep and Glutamine. HELA cells were seeded in 6-well plates at 150000 cells per well two days prior to Lentiviral infection. After two days, 2.5 ml of viral supernatant, supplemented with protamine sulphate, was added per well. Plates were incubated for two days. After two days the medium was replaced with DMEM complete + 5 ug/ml blasticidin for selection. The following day cells were passaged into 10 cm culture dishes and maintained in DMEM complete + 5 ug/ml blasticidin.

### IP-MS

Cell lines expressing the respective V5-tagged protein (V5-JTB and V5-SYS1) were grown in 4x10 cm culture dishes until confluent. Cells were harvested by scraping in cold PBS-EDTA, washed once in PBS without EDTA and the pellet was snap-frozen in liquid nitrogen. In the morning of the IP experiments, pellets were thawed in a 37°C water bath and resuspended in 1ml lysis buffer (25 mM HEPES pH 7.5, 150 mM NaCl, 0.5% NP40 substitute, 1 tablet of EDTA-free protease inhibitor per 25 ml of lysis buffer). Lysates were incubated rotating at 4 °C for 20 minutes and then centrifuged at 17000xrcf for 20 minutes at 4 °C. 50 ul of ChromoTek V5-Trap® Agarose was washed three times with 500 ul lysis buffer; spinning at 2500 x rcf to recover the beads. The supernatant of the cell lysate was added to the beads and the samples were incubated, rotating, at 4 °C for 4 hours. After incubation the beads were washed twice with 500 ul lysis buffer, then twice with 500 ul wash buffer 1 (25 mM HEPES pH 7.5, 300 mM NaCl, 0.5 % NP-40 substitute) and finally three times with wash buffer 2 (25 mM HEPES pH 7.5, 300 mM NaCl, 0.5 % NP-40 substitute). After the final wash, the supernatant was removed entirely and the beads were flash-frozen in liquid nitrogen and stored at -80 °C.

### Proteomics

Proteins were reduced and alkylated by heating immunoprecipitation beads for 7 min. at 95^0^C in 1x S-Trap Lysis buffer (5% SDS, 50 mM TEAB pH 8.5) supplemented with 5 mM TCEP and 20 mM chloroacetamide. After loading TFA-acidified samples on S-Trap Micro spin columns (Protifi, NY, USA) and washing with TEAB/methanol wash buffer according to the manufacturer’s instructions, proteins were digested for 2 hrs at 47^0^C with trypsin (Sigma-Aldrich; 2µg per sample). Peptides were eluted, vacuum-dried and stored at -80°C until LC-MS/MS analysis.

Peptides were analyzed by nanoLC-MS/MS on an Orbitrap Exploris 480 Mass Spectrometer connected to an Evosep One LC system (Evosep Biotechnology, Odense, Denmark). Prior to LC separation, peptides were reconstituted in 0.1% formic acid and 10% of the sample was loaded on Evotip Pure™ (Evosep) tips. Solvent A was 0.1% formic acid in mass-spec grade water and solvent B was 0.1% formic acid in 100% ACN. Peptides were eluted directly on-column and separated using the pre-programmed “Extended Method” (88 min gradient) on an EV1137 (Evosep) column with an EV1086 (Evosep) emitter. Nanospray was achieved using the Easy-Spray NG Ion Source (Thermo Scientific) at 2000V. The Exploris 480 was operated in DDA Cycle Time mode with 1 sec. duty cycle and full MS scans being collected in the Orbitrap analyzer with 60,000 resolution at m/z 200 over a 375-1500 m/z range. Default charge state was set to 2+, the AGC target was set to “standard” for both MS1 and ddMS2 and for both scan types the maximum injection time mode was set to “auto”. Monoisotopic peak determination was set to “peptide” and dynamic exclusion was set to 20 sec. For ddMS2, a normalized HCD collision energy of 30% was applied to precursors with a 2+ to 6+ charge state meeting a 5e4 intensity threshold filter criterium. Precursors were isolated in the Quadrupole analyzer with a 1.2 m/z isolation window and MS2 spectra were acquired at 15,000 resolution in the Orbitrap.

Mass spectrometry data (RAW files) were analyzed by label-free quantitation (LFQ) with Proteome Discoverer (PD, Thermo Scientific, version 3.1.1.93) using Percolator, Minora Feature Detector and standard settings. MS/MS data were searched against the human Swissprot database (20,421 reviewed entries, release 2025_02) with Sequest HT. The maximum allowed precursor mass tolerance was 20 ppm and 0.02 Da for fragment ion masses. False discovery rates for peptide and protein identification were set to 1% and PSMs were additionally filtered for Sequest HT Xcorr score > 1. Trypsin was chosen as cleavage specificity, allowing two missed cleavages. Carbamidomethylation (C) was set as fixed modification, whereas oxidation (M) and acetyl (Protein N-term) were used as variable modifications. A maximum retention time shift of 10 min. was allowed in the consensus workflow Feature Mapper node and normalization was performed on total peptide amount, without scaling, in the Precursor Ions Quantifier node. Summed abundances were used for protein abundance calculation. Protein abundances of the .pDResult file were exported and further processed in Perseus ^52^ (version 2.1.4.0). Missing values were replaced by imputation based on random drawing from a normal distribution using a width of 0.3 and a downshift of 1.8. Differential proteins were determined using a Student’s t-test (minimal threshold:–log(p-value) ≥ 1.3 and fold change [x/y] ≥ 1 | [x/y] ≤ -1). Raw data will be deposited to PRIDE^53^.

### Immunofluorescence imaging

Hela cells (100000 per well) were seeded onto cover slips in 6-well plates. The following day, cells were washed three times with 1 ml PBS and then fixed in PBS + 4% paraformaldehyde (methanol-free) for 15 minutes. Fixed cells were washed three times with 1 ml PBS and then permeabilized for 3 minutes in PBS + 0.1% triton, followed by another three washes with PBS. Cells were then incubated in blocking buffer (PBS, 1% BSA, 0.1% Tween-20) for 60 minus. Antibodies against the V5 tag (abcam , ab27671) and against GOLGA5 (Sigma-Aldrich HPA000992) were diluted at a 1:500 ratio in PBS + 1% BSA + 0.1% Tween-20. After incubation, the blocking buffer was removed and the respective antibodies were added to the cells. Cells were incubated with the antibody over night at 4 °C. The antibody solution was removed after incubation and cells were washed with 1 ml PBS + 1% BSA + 0.1% Tween-20 three times. Fluorescently labelled anti-rabbit (Thermo Fisher, A-11011) and anti-mouse (Thermo Fisher, A-21202) antibodies were diluted 1:5000 and added to the cells. Cells were incubated with secondary antibodies for 90 minutes, then washed three times in 1 ml PBS and mounted onto a coverslip. Images were collected with a Zeiss Axio Observer 7 inverted widefield fluorescence microscope.

## Supplemental info

**Supplemental Figure S1:**
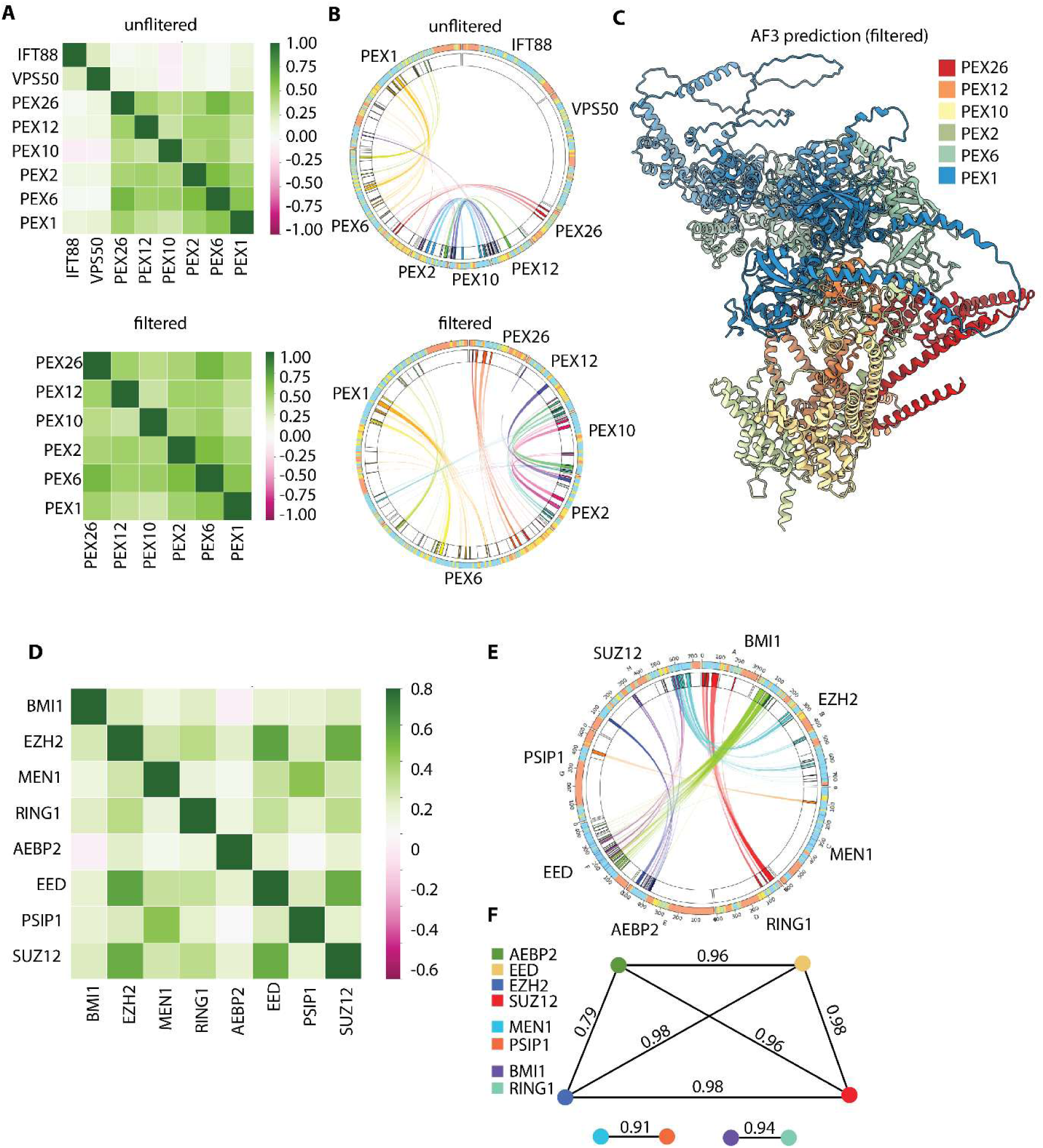
AlphaBridge filtering removes spurious interactions. **A** DepMap correlation matrices before (top) and after (bottom) AlphaBridge filtering. **B** AlphaBridge plots before (top) and after (bottom) filtering. Proteins are shown as a broken circle with connecting lines indicating confidently predicted interactions as judged by AlphaBridge metrics **C** AF3 structure prediction of the complex after filtering. **D** DepMap correlation matrix containing multiple distinct protein complexes. **E** AlphaBridge plot of an AF3 prediction for all proteins in **(D)**. Network representation of the AlphaBridge output highlighting the presence of three different protein complexes. Nodes are proteins and edges represent confident interactions with AlphaBridge scores.

**Supplemental Figure S2:**
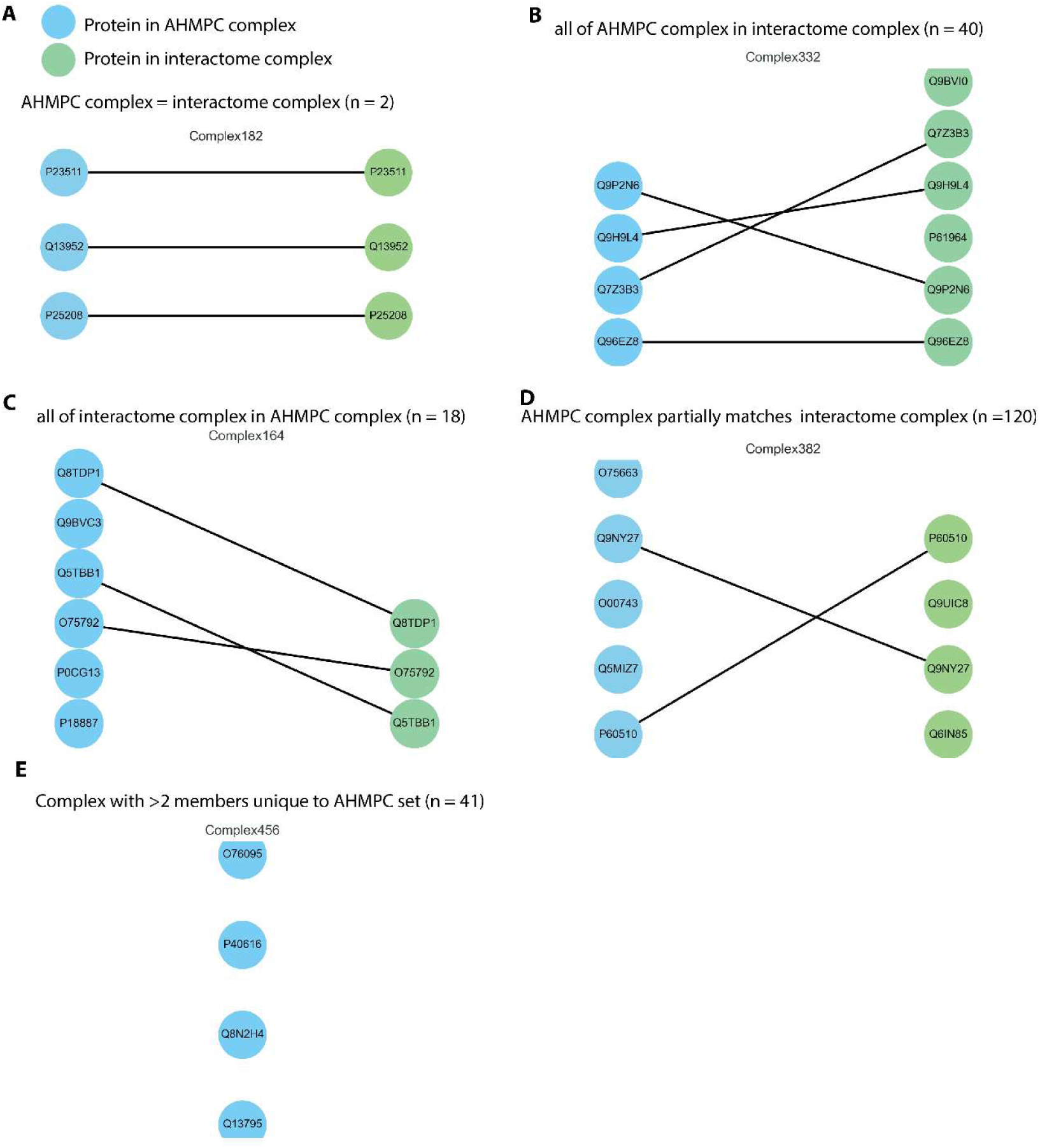
AHMPC and interactome datasets include some overlap and many distinct complexes. **A** Complete match of a AHMPC to interactome complex. Network representation of complexes showing proteins as nodes. AHMPC complexes are shown in blue and interactome complexes in green. Edges indicate where proteins are present in both complexes. Uniprot IDs are written in each node. **B** Same representation as **(A)** but illustrating a case where all members of the AHMPC complex are contained in a larger interactome complex. **C** Same representation as **(A)** but illustrating a case where all members of the interactome complex are contained in a larger AHMPC complex. **D** Same representation as **(A)** but illustrating a case where some members of the AHMPC complex are matching some members of a interactome complex. **E** Illustrating complexes that are unique to the AHMPC set.

**Supplemental Figure S3:**
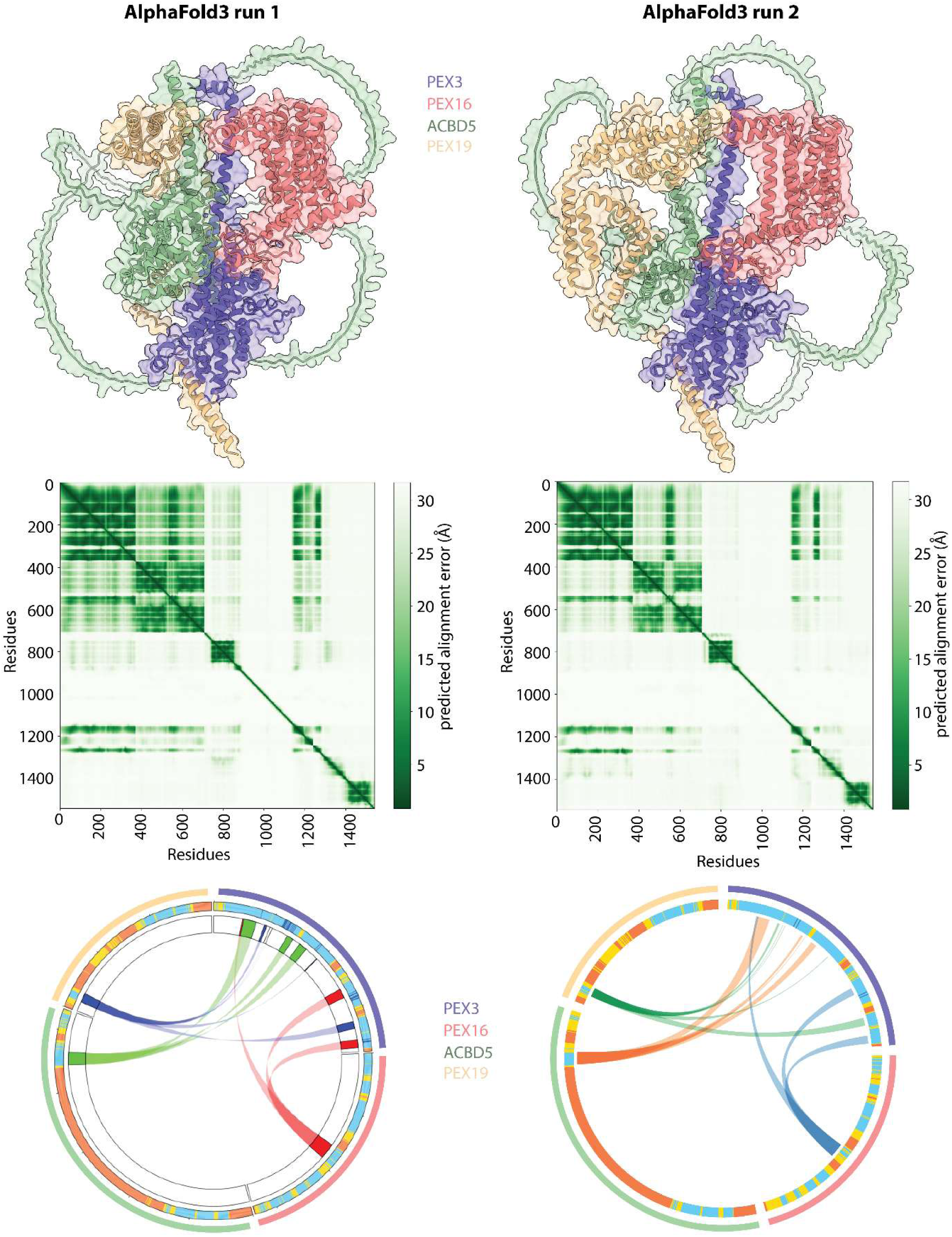
Comparison of different predictions for the same complex. While relative orientations of subunits can difers in different predictions, confidence metrics and AlphaBridge results show that predicted interaction sites are consistent.

**Supplemental Figure S4:**
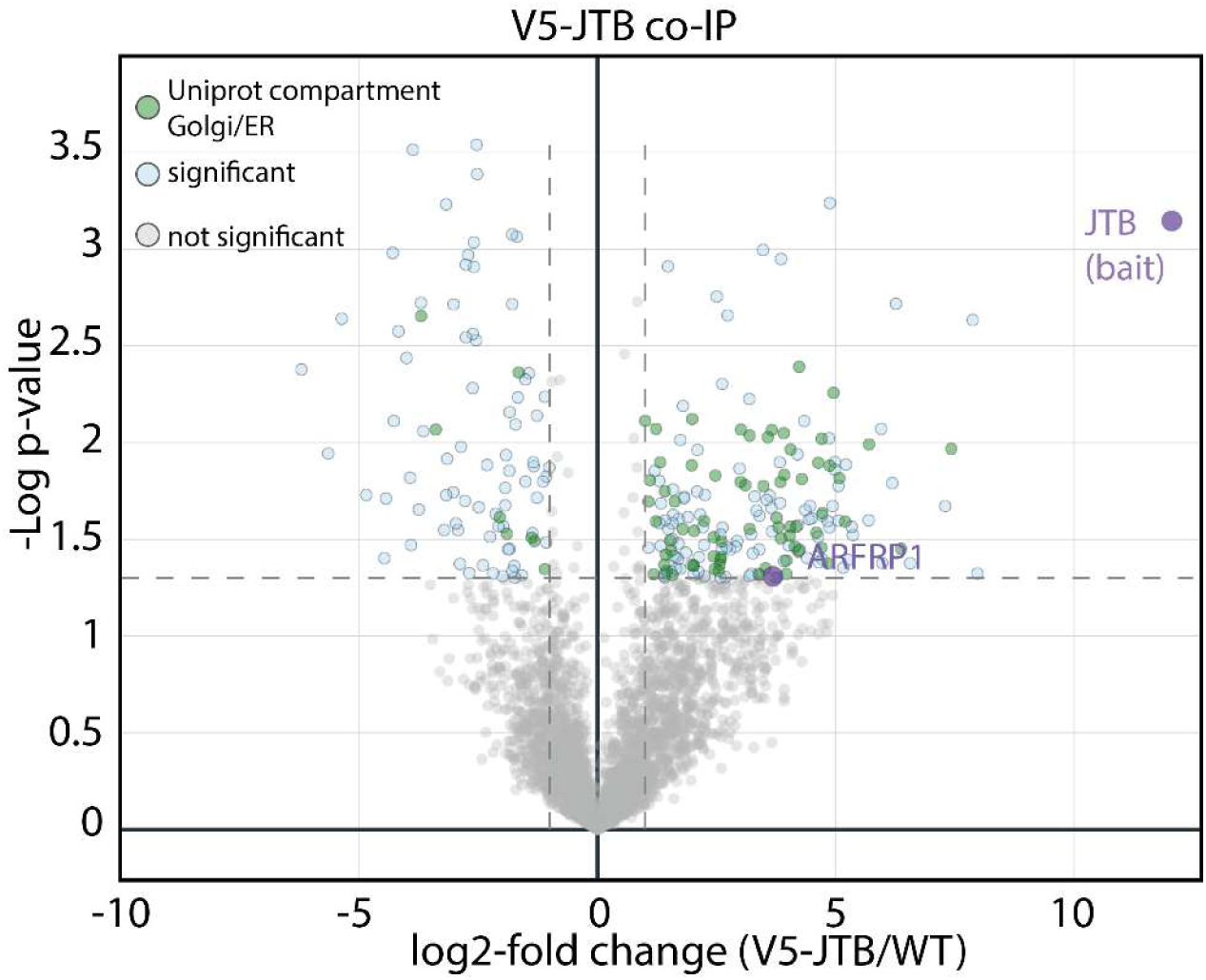
IP-MS with V5-tagged JTB. IP-MS using V5-SYS1 expressed in HELA cells as bait. Significance cutoff: Student’s T-test p-value of 0.05 calculated on three replicates. Proteins that have Uniprot compartment annotation “Golgi” or “ER” are coloured in green.

**Supplemental Figure S5:**
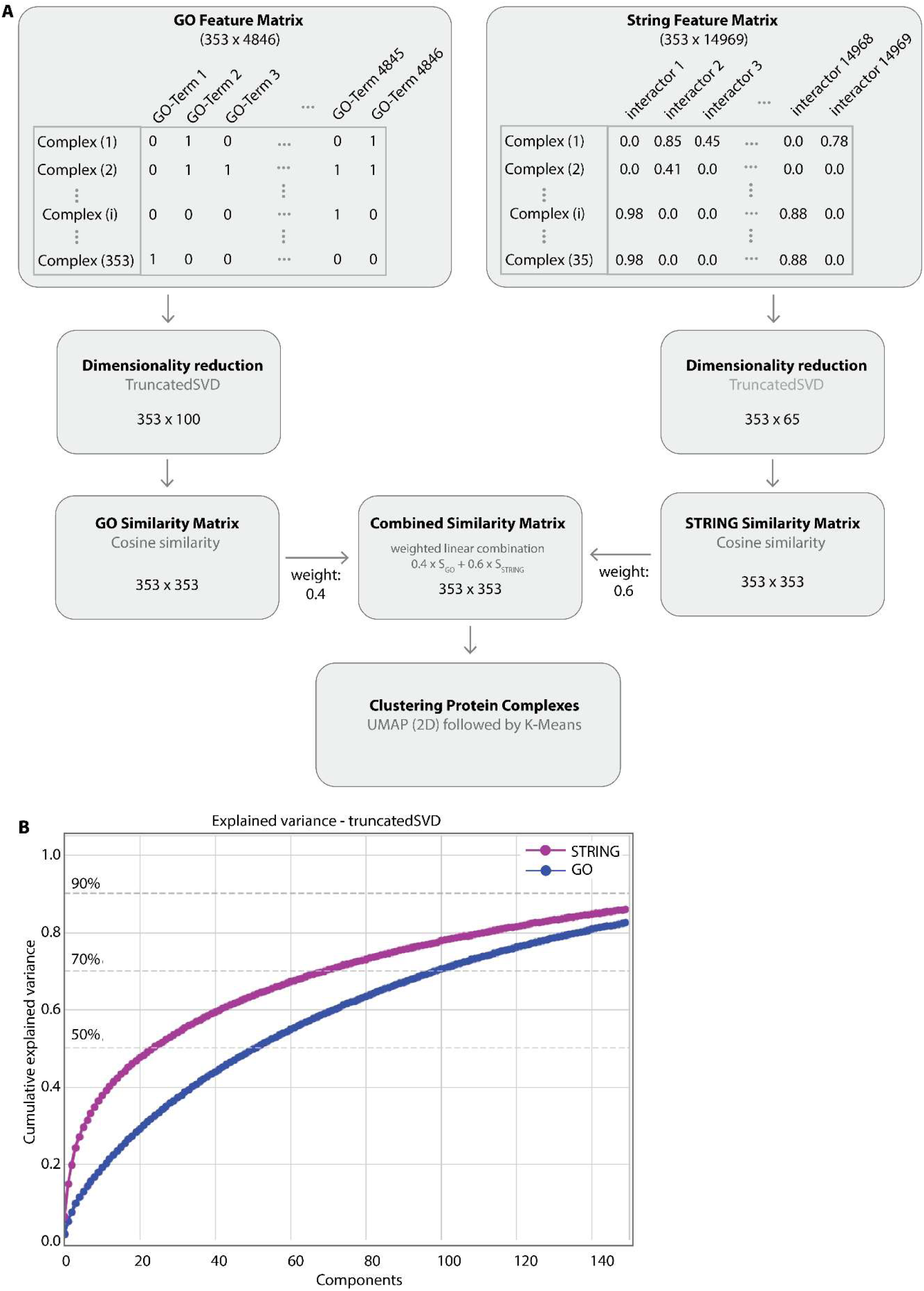
Functional embeddings. **A** Flowchart of the data processing to generate functional embeddings. The values in GO Feature Matrix and STRING feature matrix are only for illustration and do not reflect the actual data. **B** Cumulative explained variance of the truncatedSVD dimensionality reduction plotted for increasing number of components.

**Supplemental Figure S6:**
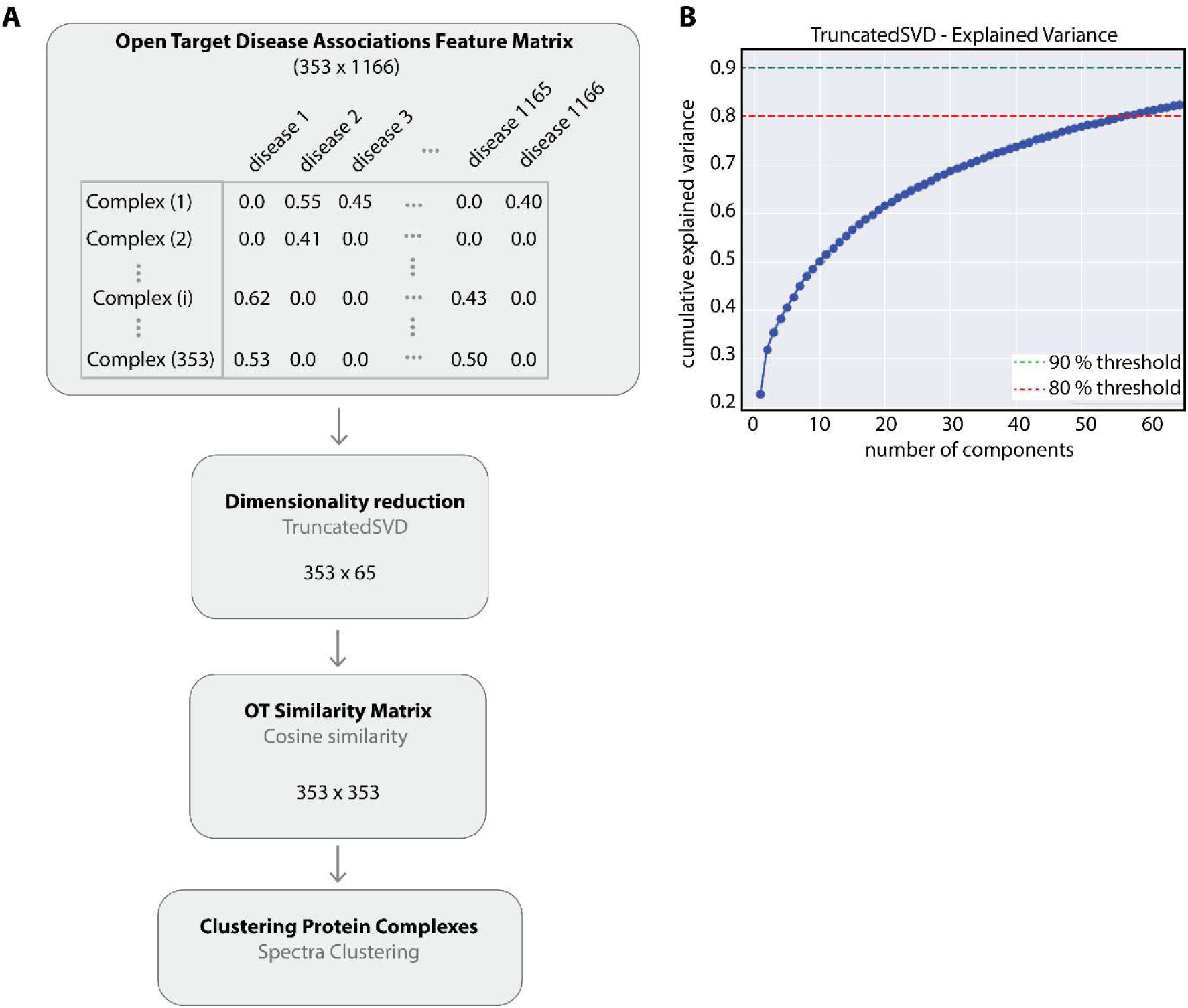
Disease associated embeddings. **A** Flowchart of the data processing to generate embedding of disease association. The values in the matrix are only for illustration and do not reflect the actual data. **B** Cumulative explained variance of the truncatedSVD dimensionality reduction plotted for increasing number of components.

**Supplemental Table1:** Contingency table for IP-MS results using V5-JTB as bait. Fisher Exact test shows Golgi/ER proteins are overrepresented in the significantly enriched fraction (P value: 5.298^-10^, Odds Ratio 2.77).

|  | Golgi or ER | Not Golgi or ER | Totals |
| --- | --- | --- | --- |
| Significantly enriched | 72 | 112 | 184 |
| Not significantly enriched | 641 | 2764 | 3405 |
| Totals | 713 | 2876 | 3589 |
| P_value $5.298341402338662 \cdot 10^{-10}$,<br>Odds Ratio 2.77 | | | |

